# Cryo-EM structure of the agonist-bound Hsp90-XAP2-AHR cytosolic complex

**DOI:** 10.1101/2022.05.17.491947

**Authors:** Jakub Gruszczyk, Loic Grandvuillemin, Josephine Lai-Kee-Him, Matteo Paloni, Christos G. Savva, Pierre Germain, Marina Grimaldi, Abdelhay Boulahtouf, Hok-Sau Kwong, Julien Bous, Aurelie Ancelin, Cherine Bechara, Alessandro Barducci, Patrick Balaguer, William Bourguet

## Abstract

**Summary:** Living organisms have developed protein sensors helping them to adapt to their environment^1^. The aryl hydrocarbon receptor (AHR) is an emblematic member of this class of proteins, and a ligand-dependent transcription factor that mediates a broad spectrum of (patho)physiological processes in response to numerous substances including pollutants, natural products and metabolites^2^. However, in the absence of high-resolution structural data, a molecular understanding of how AHR is activated by such diverse compounds is lacking. Here we present a 2.85 Å cryo-electron microscopy structure of the cytosolic complex comprising AHR bound to the ligand indirubin, the chaperone Hsp90 and the co-chaperone XAP2. The structure reveals a closed Hsp90 dimer with AHR threaded through its lumen. XAP2 directly interacts with Hsp90 and the AHR ligand-binding domain, thereby acting as a brace stabilizing the entire complex. Importantly, we provide the first experimental visualization of the AHR PAS-B domain bound to a ligand, revealing a unique organization of the ligand-binding pocket and the structural determinants of ligand-binding specificity and promiscuity of the receptor. By providing unprecedented structural details of the molecular initiating event leading to AHR activation, our study rationalizes prior biochemical data and provides a framework for future mechanistic studies and structure-guided drug design.

## Main text

Also known as the dioxin receptor, the aryl hydrocarbon receptor (AHR) is a ligand-dependent transcription factor involved in the regulation of genes governing many (patho)physiological processes including cell growth and differentiation, cell cycle and migration, apoptosis, haematopoiesis and carcinogenesis^3–6^. Importantly, activation of AHR in the gastrointestinal tract participates in signalling between the enteric microflora and our immune system^7–9^. A disruption of the intestinal homeostasis, for example, has been shown to contribute in the aetiology of many disorders, including inflammatory bowel disease, Crohn’s disease, ulcerative colitis, diabetes and obesity^9^. Moreover, AHR is one of the crucial chemosensory proteins associated with the metabolism of various xenobiotics, and sustained activation of the receptor can lead to toxic outcomes^10,11^. AHR belongs to the basic helix-loop-helix (bHLH) PER-ARNT-SIM (PAS) protein family^12^. The bHLH-PAS family, present ubiquitously in all kingdoms of life, is predominantly engaged in detection of such diverse signals like endogenous compounds, foreign chemicals, light and osmotic pressure, for example^2,13^. The numerous cognate ligands of AHR exhibit distinctive chemical structures and include various compounds like polycyclic aromatic hydrocarbons (PAHs), halogenated aromatic hydrocarbons (HAHs), polychlorinated biphenyls (PCBs), indole derivatives, alkaloids, polyphenols and various pharmaceuticals, with dioxin (2,3,7,8-tetrachlorodibenzo-*p*-dioxin or TCDD) being the archetypal exogenous ligand of AHR^10^. As such, AHR is a key sensor which integrates numerous external and endogenous chemical signals, and it is now considered a promising drug target^5^.

In the absence of ligand, AHR resides in the cytoplasm forming a complex with several other partners including heat shock protein 90 (Hsp90) and co-chaperones like X-associated protein 2 (XAP2) and p23. Upon ligand binding, AHR undergoes conformational changes leading to exposure of the nuclear localization signal (NLS) and therefore triggering a translocation of the complex into the nucleus where AHR is released and interacts with the AHR nuclear translocator (ARNT). The newly formed heterodimer binds to so-called “dioxin-response element” (DRE) DNA sequences and regulates the expression of target genes^14,15^. Previous structural work revealed how AHR and ARNT heterodimerize and bind to DREs but provided no insights into the AHR cytosolic complex organization or the structural basis of ligand-binding^16,17^. Here we report the high-resolution cryo-electron microscopy (cryo-EM) structure of the human ligand-bound AHR cytosolic complex as a snapshot of the first step of AHR activation.

### Structure determination and architecture of the Hsp90-XAP2-AHR complex

We co-expressed a fragment of human AHR (residues 1-437) in the presence of Hsp90, XAP2 and p23, and purified the Hsp90-XAP2-AHR complex (Extended Data Fig. 1; see Materials & Methods section for details). We combined the protein with indirubin and using cryo-EM we obtained a 2.85 Å reconstruction of the ligand-bound AHR cytosolic complex (Fig. 1a, Extended Data Fig. 2,3 and Extended Data Table 1). The structure reveals in its core a nucleotide-bound Hsp90 dimer (molecules denoted Hsp90A and Hsp90B, respectively) adopting a closed conformation (Fig. 1b). Hsp90 forms extensive interaction sites with the PAS-B domain of AHR and XAP2 that occupy the same side of Hsp90, reminiscent of the recently reported structure of the glucocorticoid receptor (GR) complex (Hsp90-p23-GR)^18^. The structure of the bHLH and PAS-A domains of AHR could not be unambiguously resolved during image processing, indicating high dynamics of this region.

**Fig. 1.**
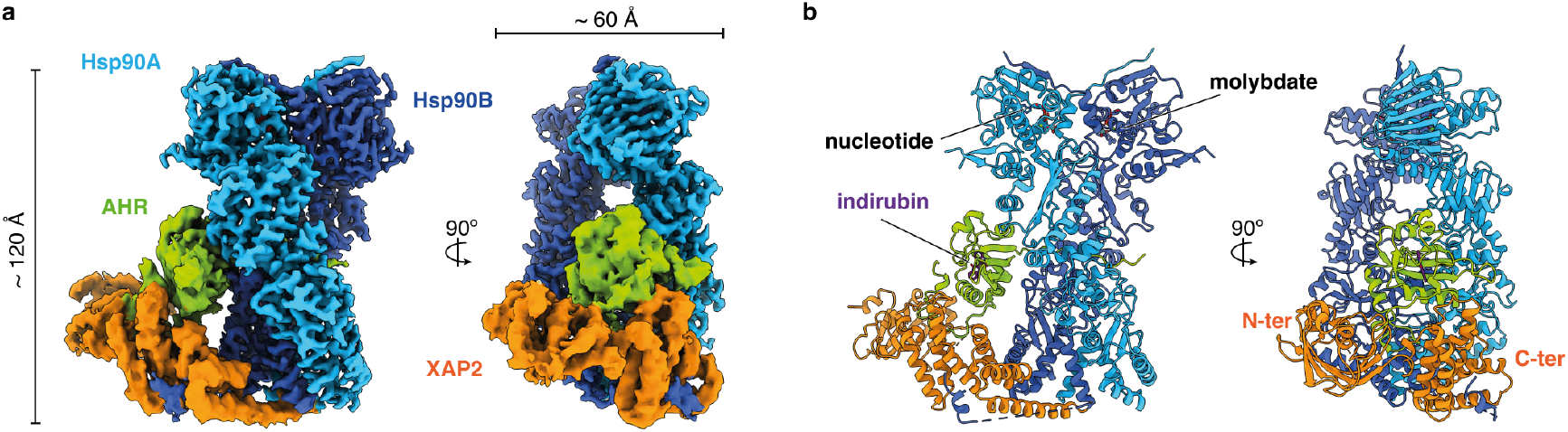
Overall architecture of the agonist-bound cytosolic complex of A HR. **a**, A composite map of the Hsp90-XAP2-AHR complex in two orthogonal views. Hsp90A (light blue), Hsp90B (dark blue), XAP2 (orange), AHR (green). The same color scheme is used throughout the manuscript unless stated otherwise. **b**, The atomic model of the complex in cartoon representation (the orientation of the molecule same as in a). Nucleotide molecule, molybdate ion and indirubin ligand are shown in sticks, The corresponding binding sites are indicated.

The PAS-B domain of AHR exhibits a canonical fold comprising a five-stranded antiparallel β-sheet (Aβ, Bβ, Gβ, Hβ, Iβ) flanked by four consecutive α-helices of variable lengths (Cα, Dα, Eα, Fα). Two additional short α-helices (Jα and Kα) are present at the carboxy-terminus of the domain (Extended Data Fig. 4a, b). The PAS-B domain is surrounded by additional extensions at both ends. The aminoterminus includes an elongated 15-residue linker region interconnecting PAS-A and PAS-B domains threaded through the Hsp90 dimer lumen (Extended Data Fig. 4a, c), and the 40 amino acid residues connecting PAS-B to the carboxy-terminal transactivation domain are forming a long loop that folds back onto the PAS-B domain and participates in the interface with XAP2 (Extended Data Fig. 4a, c).

XAP2 interacts with both Hsp90 and AHR (Fig. 1a). The amino-terminal domain of XAP2 is flexible and not well resolved in the general map. However, a focused refinement of the cryo-EM data yielded a 4.07 Å resolution map with an improved quality for the whole XAP2 protein and the carboxy-terminus of AHR (Extended Data Fig. 2,3). This is, to our knowledge, the first report of the full-length XAP2 structure.

### Hsp90 forms a core of the complex

All components of the complex are forming multiple interaction sites with each other (Fig. 2a). Interaction between AHR and Hsp90 involves two interfaces. The first interface encompasses 731 Å^2^ of buried surface area (BSA) and includes AHR residues from Aβ (I286-K290), Bβ (G299-D301), Gβ (M348), Hβ (Q364-N366) and Iβ (T382-R384) and Hsp90 residues W312, R338, A339, P340 and F341 from the middle domain of Hsp90A and Y596, T616, Y619 and M620 from the carboxy-terminal domain of Hsp90B (Fig. 2b). Note that residues localized in the β-strand Aβ were previously identified to be important for binding to Hsp90^19^.

**Fig. 2.**
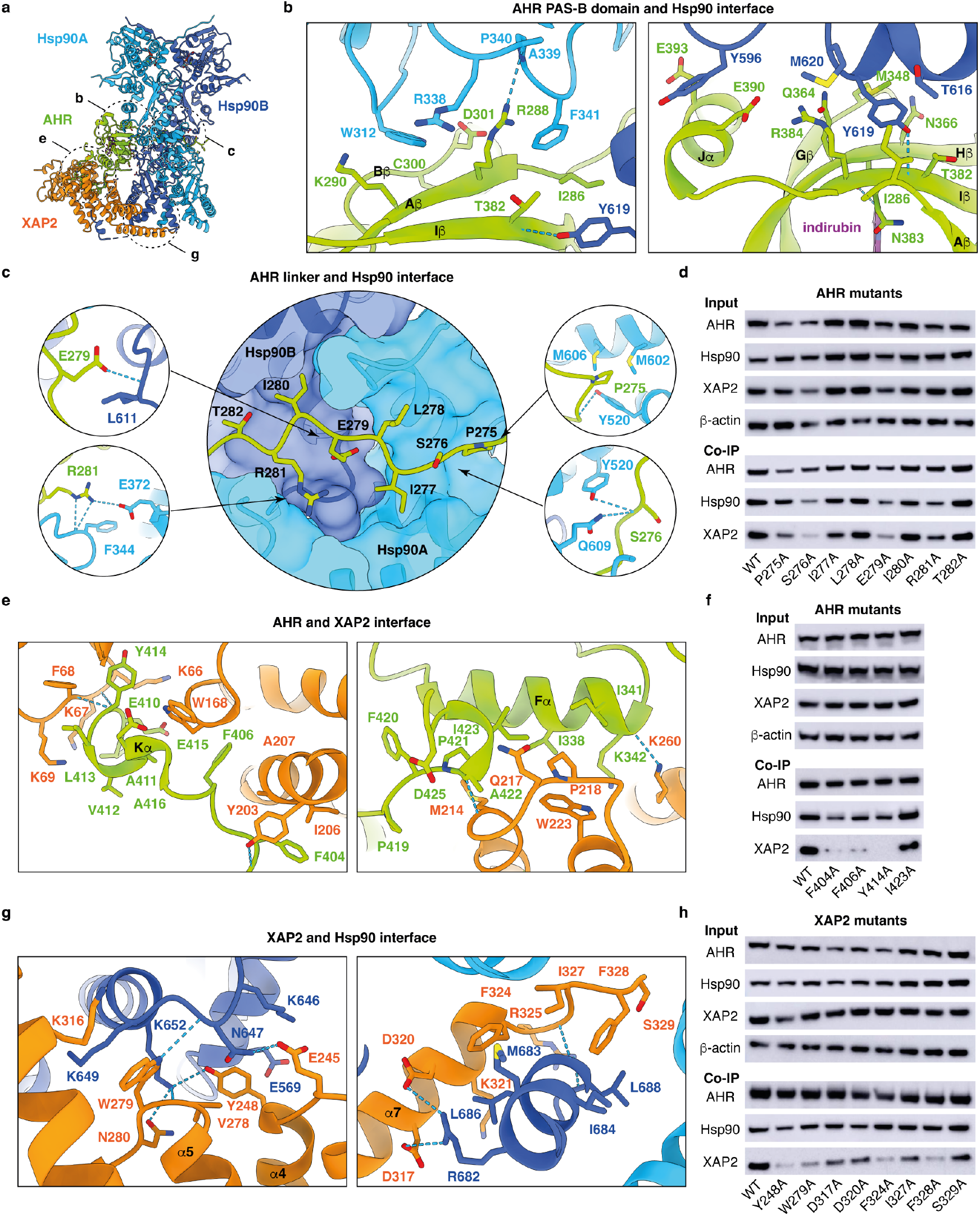
Details of the Hsp90-XAP2-AHR complex organization. **a**, An overall view of the complex with indicated location of the interaction sites between the proteins. The dashed circles indicate the contacts described in the following panels. **b**, Two principal interaction sites between Hsp90 and PAS-B domains. **c**, Close-up view of the AHR linker that is threaded through the Hsp90 lumen and interacts with both molecules. The residues for which we observed the strongest effect of the mutation on the complex stability in Co-IP assay are shown in small circles. **d**, Results of Co-IP for the mutation of the AHR linker. **e**, Close-up view of the interaction sites between AHR and XAP2. **f**, Co-IP results for the AHR mutants at the interface with XAP2. **g**, Interactions between XAP2 and Hsp90. **h**, Co-IP results for the XAP2 mutants at the interface with Hsp90. Co-IP experiments were successfully reproduced at least three times.

The second interface, confining 1,223 Å^2^ of BSA, encompasses the 15-residue linker between the PAS-A and PAS-B domains (P271-F285) that adopts an extended conformation (Fig. 2c and Extended Data Fig. 5a). The linker is threaded through the Hsp90 lumen formed by two Hsp90 molecules, thereby positioning the two PAS domains on the opposite sides of the Hsp90 dimer (Extended Data Fig. 4c). The middle and carboxy-terminal domains from both Hsp90 chains contribute to this interface by providing a combination of hydrophobic and hydrophilic residues that are engaged in multiple contacts with the backbone and side chains of AHR (Fig. 2c and Extended Data Fig. 5a). We observed that AHR I277 and I280 occupy two small hydrophobic pockets within the Hsp90 lumen, similar to L525 and L528 in the Hsp90-p23-GR^18^ and to V89 and V92 in the Hsp90-Cdc37-Cdk4^20^ structures (Extended Data Fig. 5b, c). Hence, the aliphatic nature of residues at these positions might be a hallmark for client protein interaction with Hsp90. To validate these interactions, we mutated AHR residues P275 to T282 into alanine, and monitored the impact of each mutation on AHR-Hsp90 interaction using co-immunoprecipitation (Co-IP) assays. The results revealed that the most drastic effects are obtained for the mutation of polar/charged residues (S276, E279, R281), whereas the replacement of aliphatic residues (*e*.*g*., I277, L278 and I280) by alanine has little or no impact on the stability of the complex (Fig. 2d). These data suggest that aliphatic residues at positions 277 and 280 are important for AHR recognition by Hsp90, whereas polar residues play a major role in the stability of the interaction.

### Sodium molybdate mimics ATP γ-phosphate

Sodium molybdate is known to stabilize the client-Hsp90 interaction and to inhibit gene regulation by several nuclear receptors, including AHR^21^, but its mode of action has remained elusive. Our map displays an unambiguous density for the nucleotide binding site in both Hsp90 chains (Extended Data Fig. 6a). We observed an unusually strong density at the expected position of ATP γ-phosphate, much stronger than either α- or β-phosphate (Extended Data Fig. 6b). We reasoned that, rather than the ATP γ-phosphate, this density could be that of a molybdate ion that was used in the buffer for complex preparation. We confirmed the presence of the molybdate inside the Hsp90 complex using mass spectrometry (Extended Data Fig. 6b). Therefore, we concluded that the nucleotide binding site contains one ADP molecule generated from ATP hydrolysis, one magnesium ion, and one molybdate ion that is also involved in the chelation of the magnesium ion (Extended Data Fig. 6c). Hence, by mimicking the γ - phosphate upon ATP hydrolysis, the molybdate ion would hold the Hsp90 dimer in the ATP-induced closed state, thereby preventing AHR release^22^ (Extended Data Fig. 6d).

### XAP2 stabilizes the complex

The XAP2 protein is composed of an amino-terminal FKBP-type peptidyl-prolyl cis/trans isomerase (PPIase) domain and a carboxy-terminal tetratricopeptide repeat (TPR) helical domain^23,24^ (Extended Data Fig. 7a). Our structure reveals the molecular basis of XAP2 interaction with AHR and Hsp90 suggesting that the co-chaperone serves as a brace interconnecting the other components of the complex (Fig. 1). The PPIase and TPR domains are connected by a 12-residue long hinge and adopt an open conformation without any direct inter-domain interaction (Extended Data Fig. 7a). The interaction between AHR and XAP2 (1,165 Å^2^) is mainly conferred through helix Fα, while the XAP2 regions in contact with AHR include the loop between strand βC’ and helix aII (K66-K69) within the PPIase domain, helix α2 and the loop α2-α3 of the TPR domain (Y203, I206, A207, M214-P218, W223), and the linker region (W168) (Fig. 2e). Single amino acid substitutions of three AHR residues at the interface with XAP2 (F404A, F406A, Y414A) had a deleterious effect on XAP2 binding and led to reduction in Hsp90 interaction, suggesting that XAP2 helps stabilize the whole complex (Fig. 2f).

XAP2 also forms an extensive contact with Hsp90 (1,027 Å^2^) mainly via the carboxy-terminal domain of the Hsp90B and the carboxy-terminal MEEVD motif, the latter previously described^24^ (Fig. 2g and Extended Data Fig. 7b, c). We mutated XAP2 residues at the interface with Hsp90 (Y248A in α4, W279A in loop α5-α6, or D317A, D320A, F324A, I327A, F328A, S329A in helix α7). All mutations except S329A abrogated or strongly reduced XAP2 binding and decreased the interaction between AHR and Hsp90 (Fig. 2h). Together, these data suggest that Hsp90 and XAP2 are interacting with AHR in a concerted fashion, XAP2 serving as a scaffolding protein that stabilizes the cytosolic complex.

### Exploring AHR ligand-binding mechanism

Indirubin (Fig. 3a) is a dietary-derived endogenous ligand of AHR produced from tryptophan by intestinal bacteria^25^. We confirmed that indirubin is a potent activator of AHR using cell-based assay (Fig. 3b). We also demonstrated that the purified Hsp90-XAP2-AHR complex is active and binds indirubin, and that the presence of p23 is not required for the interaction with the ligand (Fig. 3c). In the cryo-EM structure, the indirubin binding site is unambiguously resolved within the PAS-B pocket (Fig. 3d).

**Fig. 3.**
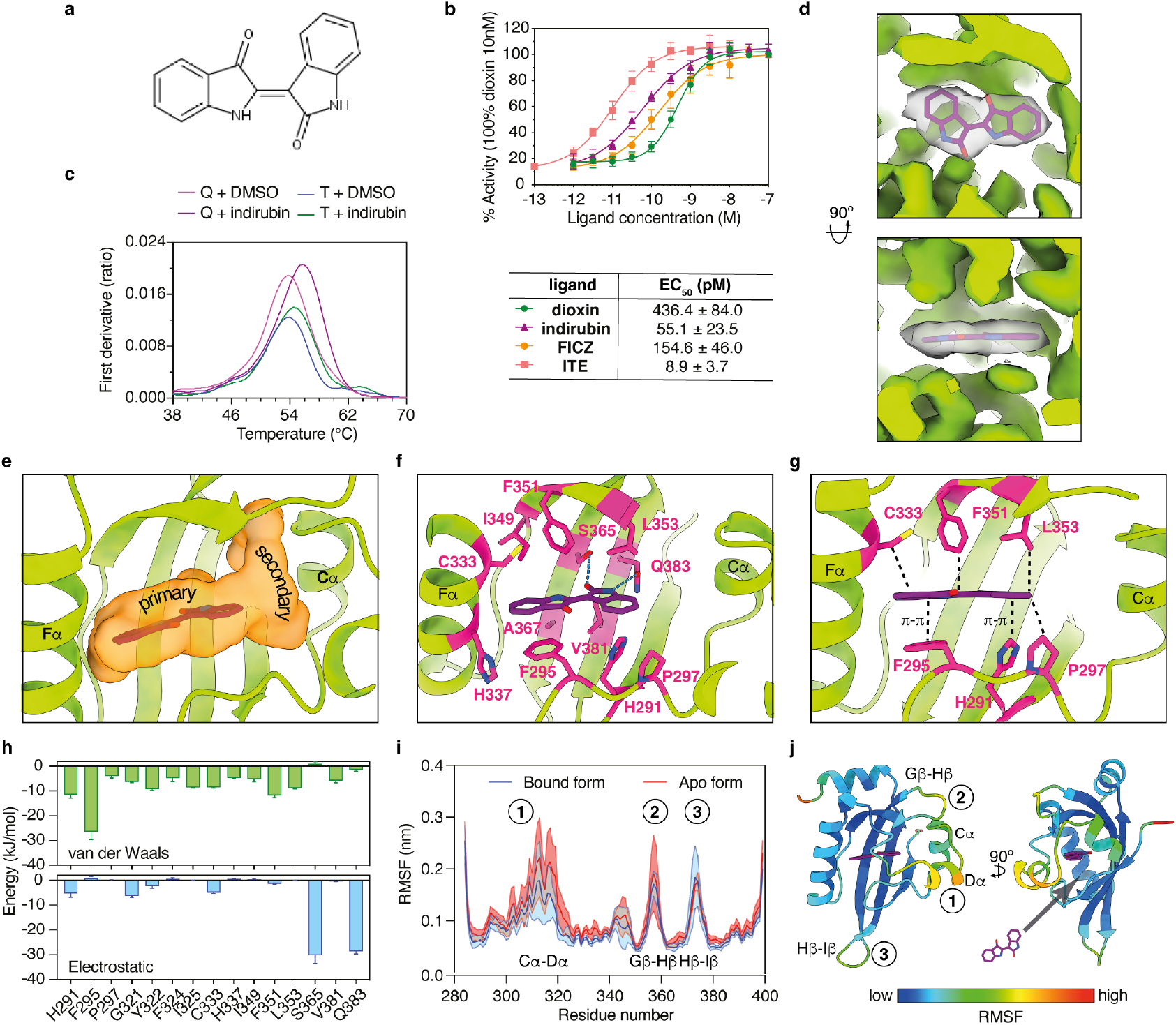
Characterization of the interaction with indirubin. **a**, Structure of indirubin. **b**, Results of cell-based activity assays for four AHR ligands. The results are representative of independent biological replicates (n=4). **c**, Two orthogonal views of the cryo-EM map for the region corresponding to indirubin. **d**, Nano-DSF analysis of the interaction between Hsp90-XAP2-AHR (T, ternary) and Hsp90-XAP2-p23-AHR (Q, quaternary) complexes and indirubin. The results are representative of independent biological replicates (n=3). **e**, Close-up view of the ligand binding pocket (LBP) shown as an orange surface. **f**, Close-up view of the indirubin binding site. Residues interacting with the ligand are shown as magenta sticks. Hydrogen bonds are indicated as dashed blue lines. **g**, Two layers of LBP residues determine AHR ligand selectivity. **h**, Van der Waals (upper panel) and electrostatic (lower panel) interaction energies between indirubin and the residues of the protein. Values represent averages between 8 MD simulations; error bars are standard deviation between the replicas. **i**, Root mean square fluctuations (RMSF) of the protein backbone in bound (blue) and apo (red) conformations. Values are averages over the MD simulations; shaded areas represent the standard deviation between the replicas. **j**, Two views of the PAS-B domain colored according to the RMSF. The location of a potential ligand entry site to LBP is indicated.

The AHR ligand-binding pocket (LBP) takes a form of an elongated channel extending perpendicularly between helices Cα and Fα (Fig. 3e). Remarkably, all elements of the PAS-B secondary structure contribute to the LBP. The amino acids lining the cavity are essentially hydrophobic (71%), including 8 aromatic (26%) and 14 aliphatic (45%) residues. The LBP also comprises 9 polar amino acids (29%) but is essentially devoid of charged residues (Extended Data Fig. 8a). The ligand occupies a volume of approximately 220 Å^3^ out of the total 682 Å^3^ of the void volume as determined by CASTp^26^. Accordingly, among 31 residues contributing the LBP, only 15 side chains interact with indirubin (4.2 Å distance cut-off), indicating that a significant portion of the cavity remains unoccupied (Fig. 3e and Extended Data Fig. 8b). The amino acids involved in binding are located in the helix Fα region and involves residues originating mainly from Bβ, Eα, Fα, Gβ, Hβ and Iβ, whereas on the other side of the pocket, the volume delineated by Ab, Ca, Da and the loop Gβ-Hβ remains unoccupied (Fig. 3f). Indirubin is a planar molecule with an asymmetric double indole structure. The indole ring adjacent to helix Fα is engaged in many contacts with aliphatic and aromatic pocket residues, namely F295, Y322, I325, C333, H337, I349 and F351. The second indole ring is primarily involved in two hydrogen bonds with S365 and Q383 via its carbonyl and amine moieties, respectively, in addition to several van der Waals contacts with H291, P297, G321, F324, F351, L353 and V381. The hallmark of most AHR ligands is their planarity. Our structure reveals that indirubin intercalates between H291, F295 and P297 on one side of the ligand and I325, C333, F351 and L353 on the other side, the two layers of residues exerting a critical role in the selection of indirubin, and most likely other AHR ligands, based on its planar structure (Fig. 3g).

Multiple microsecond-long molecular dynamics (MD) simulations were carried out to obtain an estimation of the relative energetic contributions of the LBP amino acids to the interaction between indirubin and the PAS-B domain. The ligand remains stable in the experimental pose and its presence stabilizes the bound conformation of the protein (Extended Data Fig. 9a), maintaining stable contacts with most of the residues observed in the cryo-EM structure (Extended Data Fig. 9b). Most of these amino acids form favourable Lennard-Jones interactions with indirubin (Fig. 3h, upper panel), in particular three residues with aromatic side chains form stable stacked or T-shaped π-interactions with the indole rings of indirubin, namely H291, F295, and Y322 (Fig. 3g and Extended Data Fig. 9c, d). Our MD simulations confirms the substantial implication of S365 and Q383 that are involved in stable hydrogen bonds with the ligand (Fig. 3h, lower panel). Interestingly, the presence of indirubin reduces the overall mobility of the backbone, in particular helices Cα, Dα, and the loop Gβ-Hβ (Fig. 3i, j). The high dynamics of these two neighbouring regions in the AHR apo form suggests a possible entry site for the ligand.

To experimentally validate the indirubin binding mode, we next mutated several interacting residues and tested the binding properties of the purified mutant proteins using nano-DSF (Extended Data Fig. 10a). We observed that substitution of H291, F295 and S365 by alanine residue strongly reduced interaction with indirubin. Previous reports revealed significant species differences in AHR response to indirubin treatment^27^. Accordingly, we found that indirubin activates the human receptor more potently than the rat and zebrafish homologues (Extended Data Fig. 10b). In full support of these data, sequence (Extended Data Fig. 10c) and structural (Extended Data Fig. 10d) analyses revealed that important stabilizing interactions provided by I349 and V381 in human AHR are lost in the two other species which contain smaller residues at these positions (T and A, respectively). In the fish receptor, additional replacement of human S365 by an alanine residue is also very likely to explain a decrease in indirubin response due to removal of a stabilizing hydrogen bond with the ligand.

Together, these observations support the notion that the indirubin binding site adjacent to helix Fα is the primary anchor point of AHR ligands (Fig. 3e). This region of the LBP harbours all structural and molecular determinants controlling ligand-binding specificity, promiscuity and affinity by imposing strong geometrical constraints in order to fit flat hydrophobic ligands (*e*.*g*., PAHs, HAHs, etc.), whilst providing polar amino acid residues with the potential to form hydrogen bonds with hydrophilic chemical groups (*e*.*g*., indole derivatives such as indirubin). Moreover, our cryo-EM structure implies how AHR can accommodate also larger compounds. The binding pocket extension towards helix Cα, referred to as the secondary binding site (Fig. 3e), appears to be less geometrically constrained and contains a mix of aliphatic and polar side chains that can engage in diverse types of contacts with compounds featuring higher three-dimensional hindrance.

## Discussion

Attempts to express, purify and solve the structure of the AHR ligand-binding domain, either in the context of the full-length protein or as an isolated domain have failed for many years. In our approach we reasoned that the reconstitution of the cytosolic complex by co-expressing AHR with its main chaperone and co-chaperone would help structurally and functionally stabilize the receptor. Using the Sf9 insect cell expression system we were able to obtain a stable protein sample and solve the cryo-EM structure of the indirubin-bound Hsp90-XAP2-AHR complex, at a resolution of 2.85 Å. This study provides unprecedented structural insight into the details of the complex assembly. AHR is forming extensive interaction sites with both protein partners, partially explaining why all previous efforts to obtain the isolated PAS-B domain of human AHR have failed. The structure reveals how Hsp90 and XAP2 are interacting with AHR and maintaining the protein in the active form. The interactions involve not only the linker interconnecting AHR PAS-A and B domains and PAS-B itself but also the mainly disordered C-terminal extension of the protein. Our data shows that Hsp90 and XAP2 play indispensable role in the formation of the complex and our mutational analysis confirmed that both proteins are working in a concerted fashion highlighting the importance of using an entire protein complex to study the molecular mechanism behind AHR-dependent signalling pathway. Our cryo-EM structure of the complex also provides insight into a general mechanism of Hsp90-client recognition. Reminiscent of the GR and Cdk4 structures^18,20^, the AHR protein is threaded through the Hsp90 dimer lumen, suggesting that the mechanism of client recognition is conserved between different client proteins. Particularly, the position of key hydrophobic residues within the extended linker region that interact with Hsp90 seems to be conserved across all three available to date structures.

The structure explains previous observations suggesting that AHR Hsp90-binding and ligand-binding sites overlap^19^. Indeed, residues localized within PAS-B strands Aβ and Bβ are involved in the interactions with Hsp90 and the ligand. The proposed mechanism of AHR activation includes ligand binding and induction of conformational changes in AHR that could be directly sensed by Hsp90. Subsequently, the signal would be transferred to the amino-terminal part of the receptor leading to exposure of the NLS and triggering translocation to the nucleus. In case of AHR, the NLS sequence is predicted to be localized within the first 60 amino acid residues. Interestingly, as our complex was prepared in the presence of a ligand, the PAS-A domain and the N-terminal part of AHR could not be convincingly placed in the electron density suggesting high flexibility of this region. This part of the protein is therefore exposed to the solvent and can interact freely with the nuclear transport machinery including the importin complex.

One of the hallmarks of AHR is its remarkable promiscuity as the protein can bind many different ligands. Our cryo-EM structure allowed us to explain the mechanism behind selectivity towards certain types of ligands. Amino acid residues involved in the formation of the primary binding site are responsible for the selectivity towards molecules exhibiting a planar shape. Importantly, mutational analyses confirmed the importance of residues involved in the formation of π-interactions with the ligand. The structure also provides the explanation for receptor promiscuity. The ligand-binding pocket with its relatively large interior cavity can accommodate molecules of different sizes. Additionally, MD simulations provide insight into the potential entry site of the ligand. The region localized between helices Cα and Dα, exposed to the solvent and not obscured by the other components of the complex, exhibits an elevated degree of flexibility in the simulations. It is also worth noting that despite striking similarity in the overall structure, the PAS-A domain of AHR does not possess any internal cavity that could serve as a potential binding pocket. Instead, the interior of the domain is occluded by a presence of several bulky aromatic amino acid residues supporting the hypothesis that in the course of its evolution the PAS-A domain has been selected to serve primarily in protein-protein interactions.

Interestingly, previous comparison studies between human and rodent proteins also revealed intriguing interspecies differences concerning AHR specificity. Despite relatively high sequence homology between human and mouse AHR receptors (around 85%), the same chemical molecule can interact with the orthologous proteins with drastically different affinities. For example, mouse AHR binds dioxin with a tenfold higher affinity than its human counterpart^28^. In contrast, human AHR binds indirubin and indoxyl sulphate with much higher affinity compared with the mouse receptor^29,30^. Intriguingly, human AHR protein appears to have evolved towards better recognition of compounds belonging to indole family, which is most likely directly connected with its involvement in the signalling in the human digestive track^31^. Despite high promiscuity of the PAS-B domain, its specificity can still be finely tuned by small variations in the protein sequence. Indeed, our analyses of AHR polymorphism between human, rat and zebrafish combined with structural data explain the interspecies differences observed in the binding affinity for indirubin. Our structure demonstrates how microbiome-derived compound can interact with a human protein and modulate host signalling pathways. Therefore, it provides an insight into the molecular mechanism that emerged as a result of coevolution between human body and its microbiome.

By disclosing the first atomic model for the AHR cytosolic complex and for its PAS-B domain bound to a ligand, our structure has enabled us to explain many prior biochemical observations. Moreover, it provides a strong rationale for future mechanistic studies and development of novel pharmaceuticals that could find applications in the treatment of many diseases, including irritable bowel disease, allergic responses and cancer.

## Acknowledgments

We acknowledge The Midlands Regional CryoEM Facility at the Leicester Institute of Structural and Chemical Biology (LISCB), major funding from MRC (MC_PC_17136). We thank Rémi Freydier and Léa Causse (AETE-ISO platform, OSU-OREME/Université de Montpellier) for the determination of molybdate concentrations in our samples. Centre de Biologie Structurale is a member of the French Infrastructure for Integrated Structural Biology (FRISBI), a national infrastructure supported by the French National Research Agency (ANR-10-INBS-04-01). Native MS experiments were carried out at the Montpellier Proteomics Platform (PPM, Biocampus Montpellier) supported by the regional funds FEDER/Région Occitanie, MUSE and Labex EpiGenMed. The work was supported by funding from European Union’s Horizon 2020 research and innovation program grant No. 825489 (W.B.), ATIP-Avenir 2020 grant No. R20059SP (J.G.) and the ANR (the French National Research Agency) “Investissements d’avenir” programme reference No. ANR-16-IDEX-0006 (J.G.).

## Author Contributions

J.G. and W.B. initiated and supervised project. J.G. and L.G. purified protein samples. J.L.K.H. and A.A. prepared EM grids. J.G., J.L.K.H., C.G.S., J.B. and A.A. collected and analyzed EM data. J.G., H.S.K. and W.B. performed model building and refinement. J.G. and W.B. analyzed the structure. L.G. and P.G. carried out biochemical analysis. M.G., Ab.Bo. and P.B. performed cell-based assays. M.P. and Al.Ba. performed molecular dynamics simulations. C.B. collected and analyzed MS data. J.G. and W.B. principally wrote the manuscript which was finalised with input from all authors.

## Correspondence and requests for materials

should be addressed to Jakub Gruszczyk and William Bourguet.

