## Supplementary Material for "Cryo-EM structure of the agonist-bound Hsp90-XAP2-AHR cytosolic complex"

### Extended Data Figures

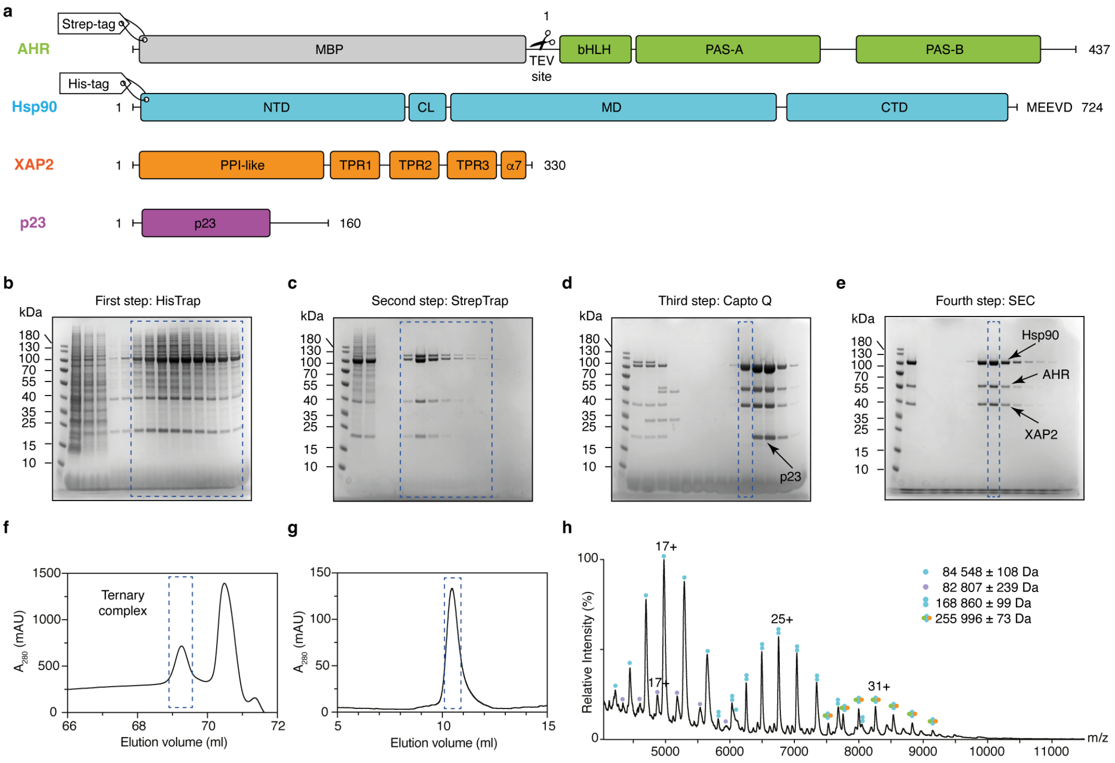


**Extended Data Fig. 1 | Preparation of the Hsp90-XAP2-AHR complex.** **a**, Schematic representation of the proteins used in this study. AHR construct includes N-terminal Strep-tag, followed by MBP protein and TEV protease cleavage site; bHLH - basic helix-loop-helix motif, PAS-A and PAS-B - Per-Arnt-Sim domain A and B. Hsp90 construct includes N-terminal His-tag; NTD - N-terminal domain, CL - charged flexible linker, MD - middle domain, CTD - C-terminal domain. XAP2 and p23 were left untagged; PPI-like - peptidyl-prolyl isomerase-like domain. TPR1, 2 and 3 - tetratricopeptide repeat motif 1, 2 and 3, α7 - C-terminal helix alpha. **b-e**, SDS-PAGE gels showing results obtained during particular steps of protein purification. Fractions used for the subsequent purification step are indicated with a blue dashed line. Molecular weight protein standards are indicated on the left-hand side of each gel. Gel images are representative of independent biological replicates (n=3). **f**, Elution profile from the ion exchange chromatography. A clear separation between ternary (no p23) and quaternary (with p23) complexes is visible; $A_{280}$ - absorbance at 280 nm, AU - arbitrary unit. **g**, Size exclusion elution profile of the Hsp90-XAP2-AHR ternary complex displaying a monodisperse peak eluting at around 10.5 ml. **h**, Native MS spectrum of the protein sample at 1.8 mg/ml, showing the presence of both monomeric (84,548 ± 108 Da, single cyan circle) and dimeric (168,860 ± 99 Da, double cyan circles) Hsp90 species as well as the intact Hsp90-XAP2-AHR ternary complexes at 255,996 ± 73 Da. Some contaminant Sf9-derived Hsp90 is also present at low intensity (82,807 ± 239 Da, light purple circle); m/z - mass-to-charge ratio. All results from protein sample purification and cryo-EM experiments were successfully reproduced at least three times. For gels source data, see Auxiliary Supplementary Information.

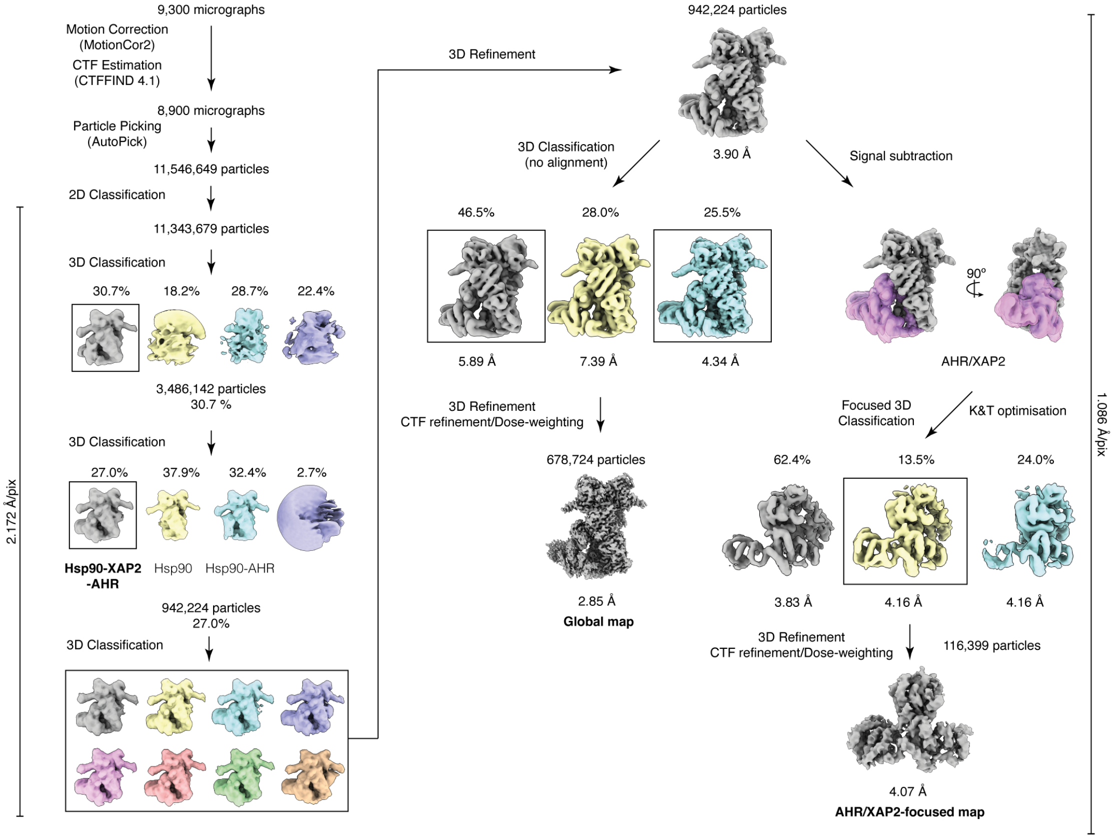


**Extended Data Fig. 2 | Cryo-EM data processing workflow.** A schematic representation of the data processing pipeline. For details, please see Materials and Methods section 'Cryo-EM data processing'. Soft mask used in focused 3D classification and refinement is shown as a semi-transparent surface and colored in purple.

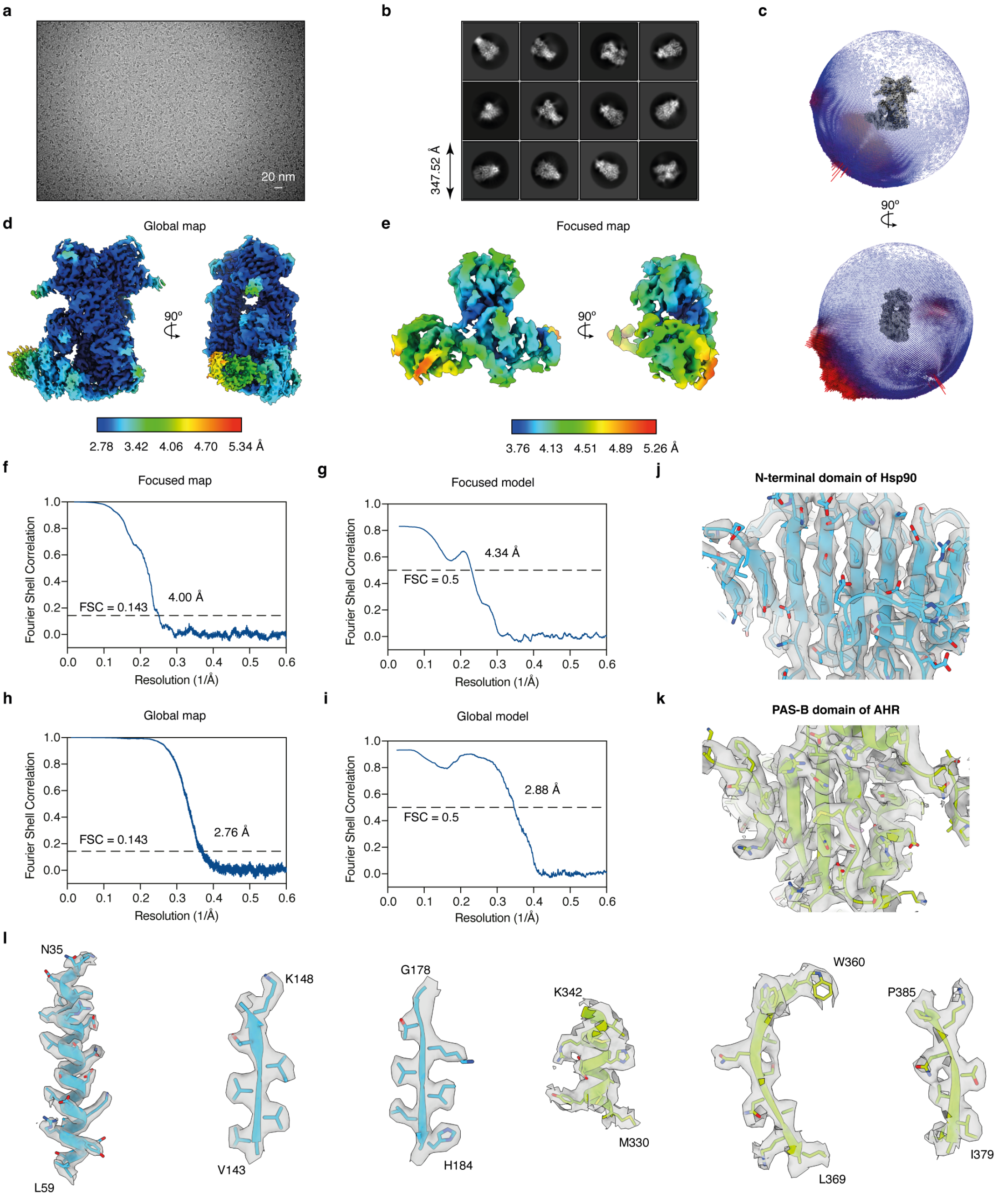


**Extended Data Fig. 3 | Details of the cryo-EM data processing. a**, Representative micrograph of the sample. The scale bar represents 20 nm. **b**, Representative 2D class averages. 2D class averages exhibit different projections corresponding to each orientation. **c**, Euler angle distribution for the final reconstruction. The model is shown in two orthogonal views. **d**, Local resolution estimation diagram of the final map - global map. Resolution key is labelled from 2.78 to 5.34 Å. **e**, Local resolution estimation diagram of the final map - focused map. Resolution key is labelled from 3.76 to 5.26 Å. **f**, Fourier-shell-correlation (FSC) plot showing the resolutions at 0.143 FSC (drawn in a dashed line) determined by gold-standard method - focused map. **g**, FSC curve of the final refined model versus the final cryo-EM map - focused model. **h**, FSC plot showing the resolutions at 0.143 FSC (drawn in a dashed line) determined by gold-standard method - global map. **i**, FSC curve of the final refined model versus the final cryo-EM map - global model. **j**, Quality of the Hsp90-XAP2-AHR cryo-EM map: N-terminal domain of Hsp90. **k**, Quality of the Hsp90-XAP2-AHR cryo-EM map: PAS-B domain of AHR. **l**, Representative examples of the electron density map for chosen elements of secondary structure.

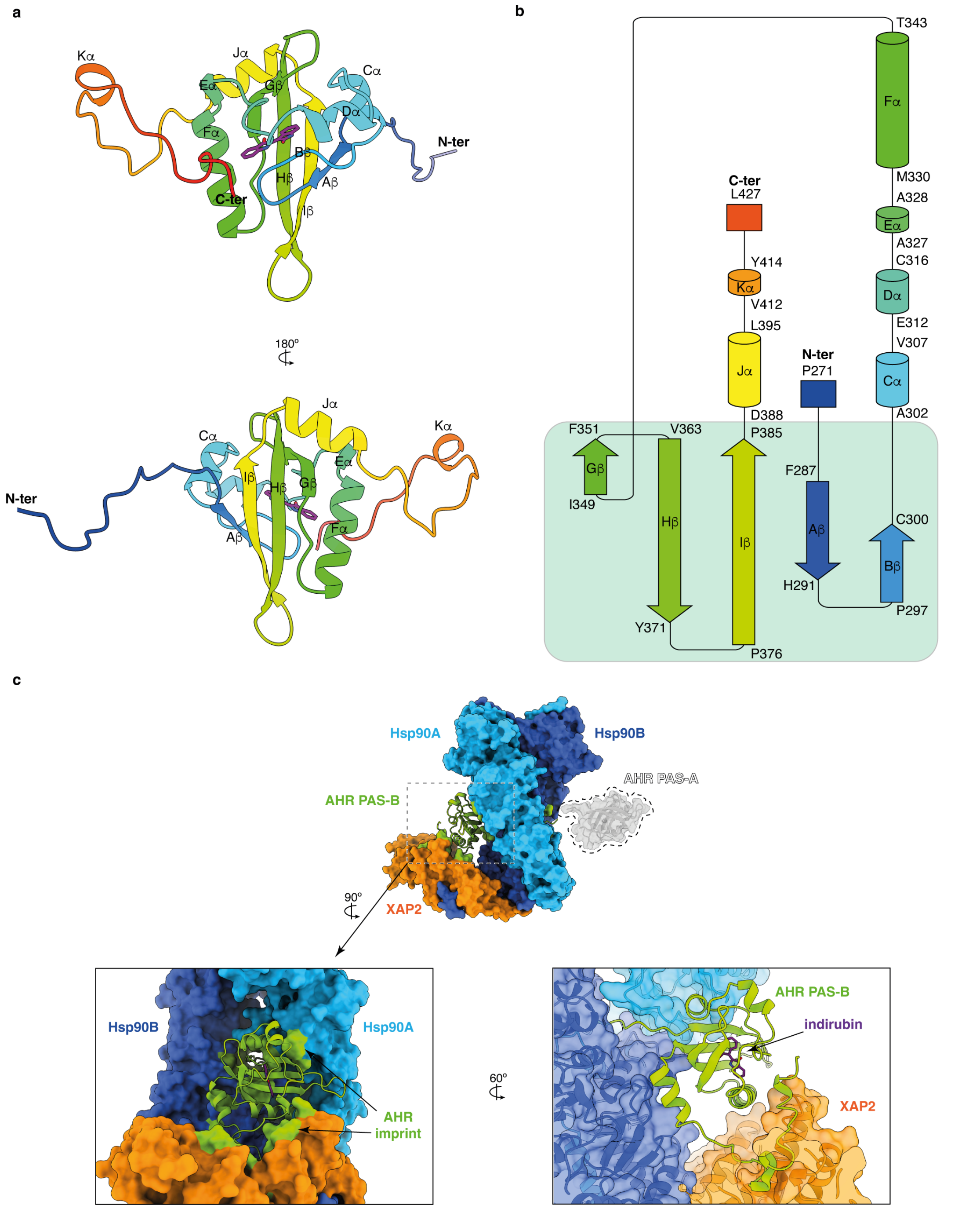


**Extended Data Fig. 4 | Organization of the AHR PAS-B domain. a**, Cartoon representation of the PAS-B domain colored in rainbow from N- to C-terminus. The elements of the secondary structure are labelled. The indirubin molecule is shown as sticks and colored in magenta. **b**, Schematic representation of the AHR PAS-B fold with indicated boundaries of the secondary structure elements. The five β-strands forming continuous β-sheet are highlighted in light green. **c**, Position of the PAS-B domain in respect to the other components of the complex. The location of the PAS-A domain (in grey) that could not be unambiguously resolved in the cryo-EM map is schematically indicated. Two close-up views show how PAS-B domain is closely surrounded by Hsp90 and XAP2. The interaction sites of the PAS-B domain with the other components of the complex are colored in green. Indirubin is shown as magenta sticks.

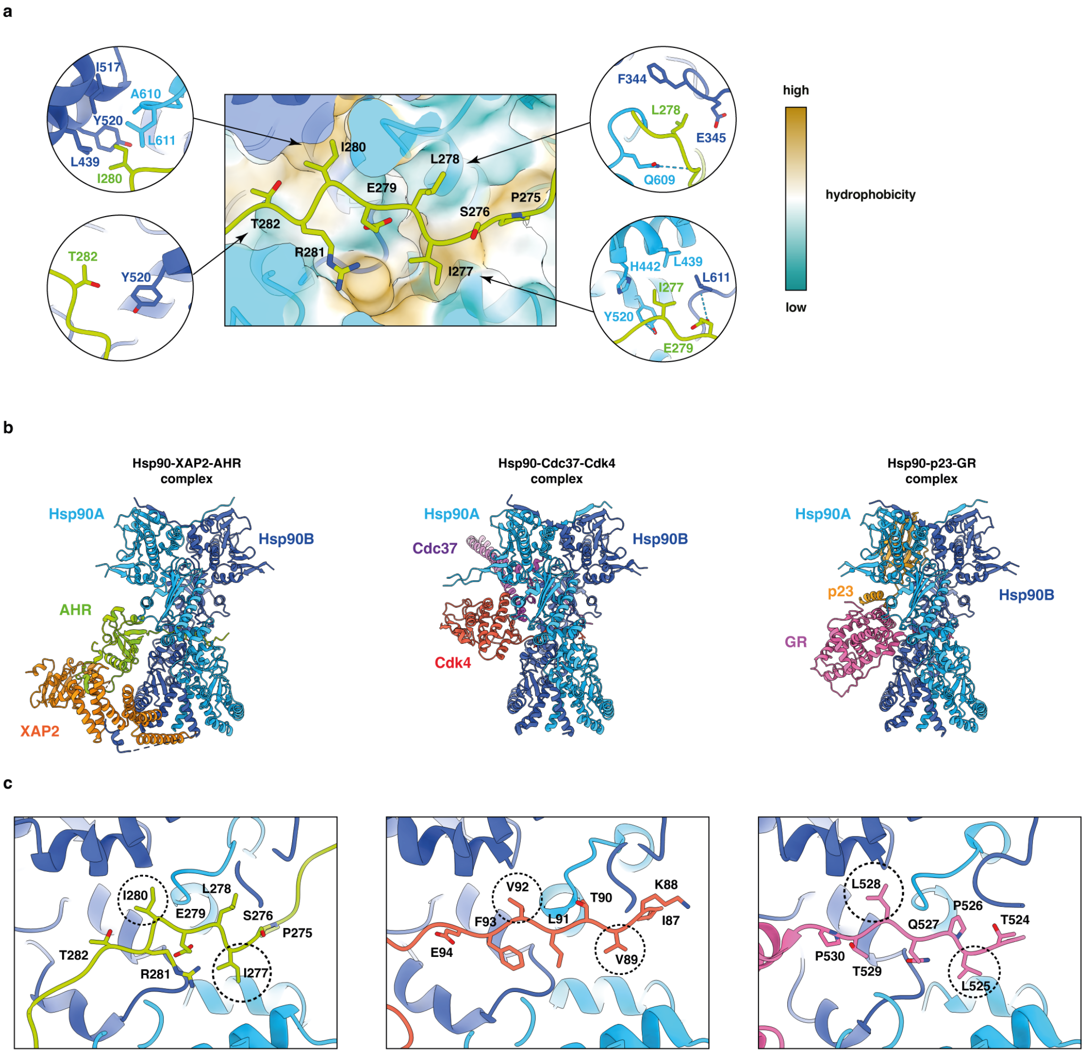


**Extended Data Fig. 5 | Detailed analysis of the Hsp90 interactions with its client proteins. a**, The AHR interdomain linker interacts with Hsp90. Two molecules of Hsp90 are shown in surface representation and colored according to the hydrophobicity of the residues. Detailed views of the interactions formed by the hydrophobic amino acid residues and T282 are shown in circles. Two isoleucine residues, I280 and I277, are facing towards hydrophobic patches located in both molecules of Hsp90. **b**, Comparison between three known structures of the Hsp90 client proteins: Hsp90-XAP2-AHR, Hsp90-Cdc37-Cdk4 and Hsp90-p23-GR complexes are shown from left to right, respectively. Hsp90 adopts a closed conformation and two molecules of Hsp90 can be superimposed with the RMSD = 0.82 Å over 1,186 aligned atoms Cα for the complexes of AHR and Cdk4 and RMSD = 0.74 over 1,217 aligned atoms Cα for the complexes of AHR and GR. The client proteins and their corresponding co-chaperons are displayed in different colors. **c**, The close-up views of the superimposition of three structures focused on the AHR, Cdk4 and GR interaction with Hsp90. In all three structures the portion of client protein interacting with Hsp90 adopts a similar extended conformation. Residues involved in the interaction are shown in sticks and labelled. The conserved positions of the hydrophobic amino acid residues are indicated with dashed circles.

a

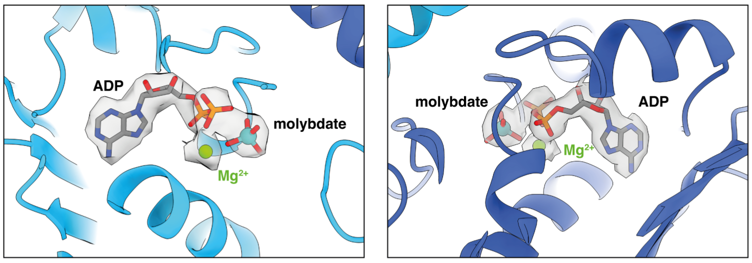


b

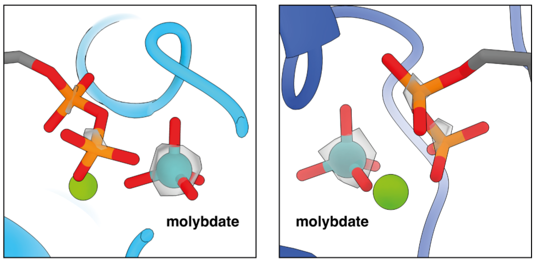


| | Sample | Molybdate concentration (μM) |
|---|---|---|
| 1 | 5 μM $Na_2MoO_4$ | 4.56 |
| 2 | 10 μM $Na_2MoO_4$ | 9.39 |
| 3 | 20 μM $Na_2MoO_4$ | 19.10 |

| | Sample | Protein concentration (μM) | Measured molybdate concentration (μM) |
|---|---|---|---|
| 1 | Protein complex | 8.3 | 8.43 |
| 2 | Protein complex + AMP-PNP | 7.1 | 0.83 |

c

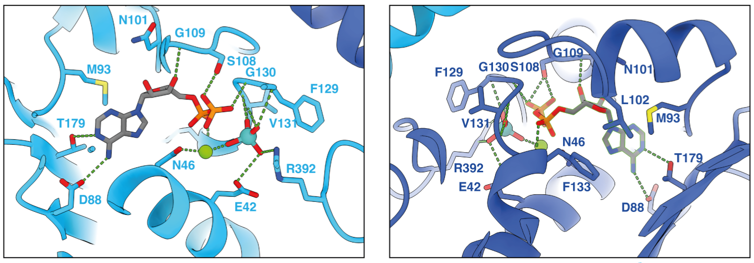


d

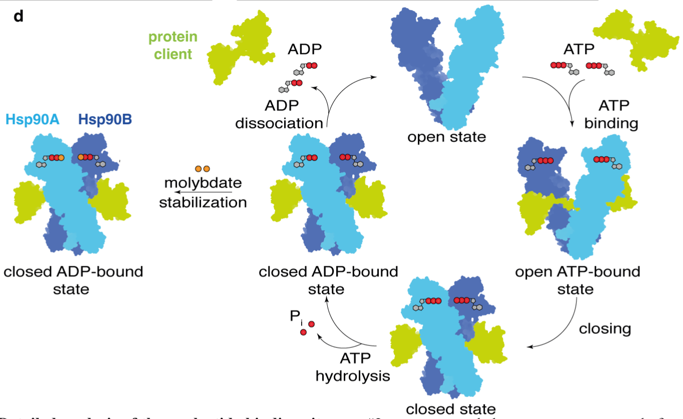


**Extended Data Fig. 6 | Detailed analysis of the nucleotide binding site of Hsp90. a**, Close-up view of the nucleotide binding site in each molecule of Hsp90. The electron density map for the molecule of ADP, molybdate anion and magnesium cation has been displayed. **b**, Close-up view of the molybdate ion. Increased contour level of the map density shows a much stronger density relative to the α and β-phosphates. Left table: Results of the analytical mass spectrometry for three standard solution of sodium molybdate at 5, 10 and 20 μM, respectively. The measured values of the molybdate concentration in the samples are in good agreement with their theoretical values. Right table: Results of the analytical mass spectrometry for two samples of the protein complex. Sample #1 was prepared as described in 'Materials and Methods' section. The measured concentration of molybdate is similar to the protein concentration in the sample. Sample #2 was prepared the same way except before buffer exchange step the protein was incubated in the presence of 5 mM AMP-PNP for 1 hour. The measured molybdate concentration in the sample is low suggesting that the non-hydrolysable analog of ATP can displace molybdate from the nucleotide binding site. The results are representative of independent biological replicates (n=2). **c**, Details of the organization of the nucleotide binding site. Amino acid residues involved in the interactions are shown as sticks and labelled. Hydrogen bonds are indicated as dashed green lines. **d**, Schematic representation of the conformational cycle of Hsp90. Binding and relaease of a client protein is coupled with binding and hydrosylis of ATP. Molybdate mimics the presence of ATP and stabilizes a closed conformation of Hsp90 even in the presence of ADP.

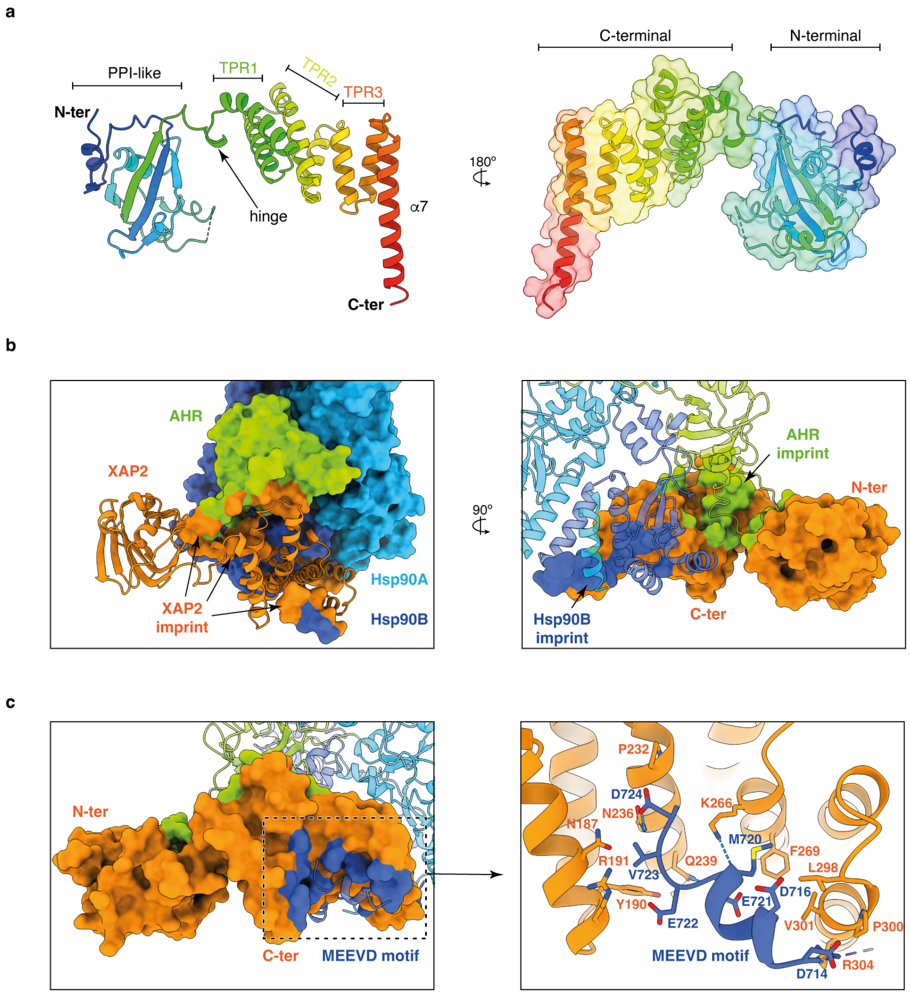


**Extended Data Fig. 7 | Detailed analysis of XAP2 protein and its interactions with the other partners. a**, Cartoon representation of XAP2 protein in two views colored in rainbow from N- to C-terminus. Location of the most important features is indicated. XAP2 is composed of two domains that are separated by a short hinge. N-terminal domain is not very well defined in the cryo-EM map most likely due to its increased mobility. The alpha-helical C-terminal domain is forming extensive contacts with both AHR and Hsp90 and is much better defined. PPI-like - peptidyl-prolyl isomerase-like domain. TPR1, 2 and 3 - tetratricopeptide repeat motif 1, 2 and 3, α7 - C-terminal helix alpha. **b**, XAP2 is interacting extensively with two other proteins. Left panel: Imprint of the XAP2 on the other components of the complex. AHR and Hsp90 are shown in surface and XAP2 in cartoon representation; the AHR and Hsp90 residues in contact with XAP2 are colored in orange. Right panel: Imprint of Hsp90 and AHR on XAP2. XAP2 is shown in surface, AHR and Hsp90 in cartoon representation. The XAP2 residues in contact with AHR and Hsp90 are colored in green and blue, respectively. **c**, Left panel: Imprint of the C-terminal part of Hsp90 on XAP2. Residues in contact with MEEVD motif are colored in dark blue. Right panel: Details of the XAP2 interactions with MEEVD motif of Hsp90. The C-terminal part of Hsp90 forms a helix that interacts with the TPR motives. Hydrogen bonds are indicated as dashed blue lines.

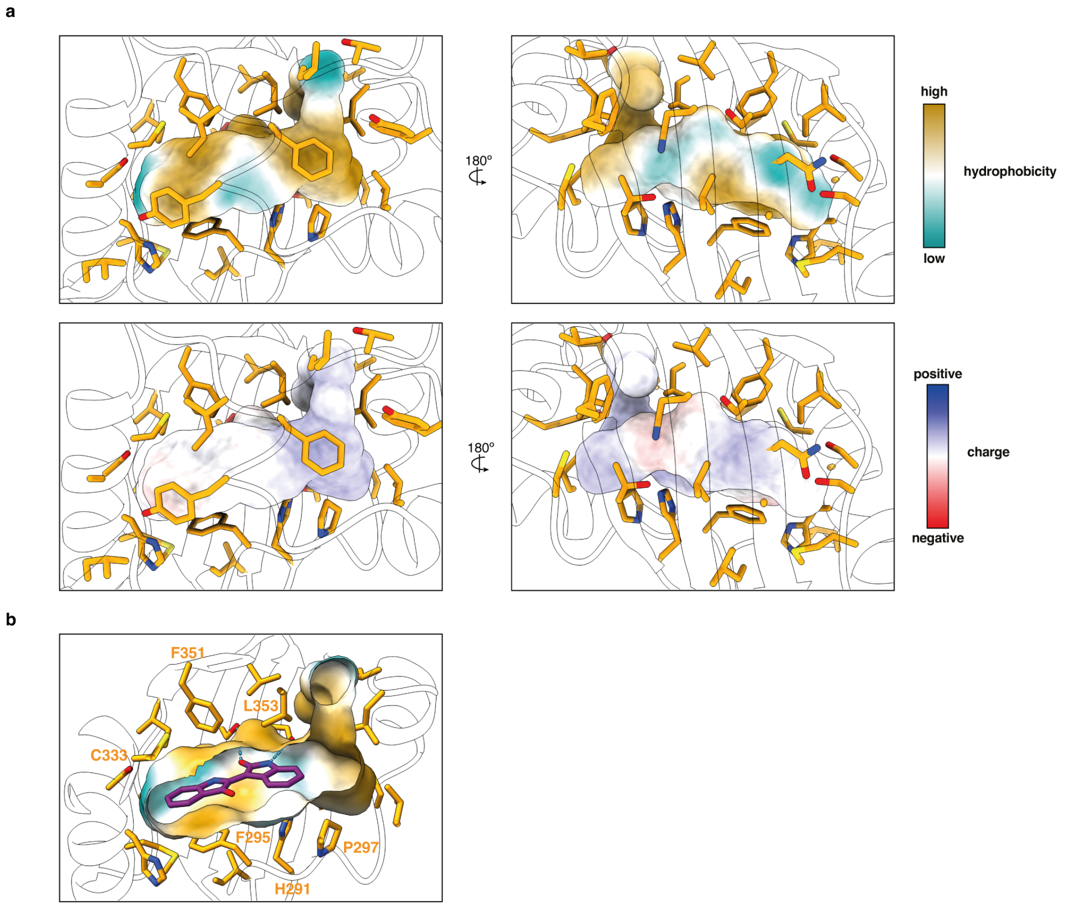


**Extended Data Fig. 8 | Detailed analysis of the ligand binding domain of AHR. a**, Amino acid residues involved in formation of ligand binding pocket (shown in sticks). The residues lining up the inside walls of the LBP are mostly hydrophobic. The surface of the LBP is colored either according to the hydrophobicity (top panel) or electrostatic potential (bottom panel). **b**, Close-up view of the interactions with indirubin. The ligand is occupying only a fraction of the LBP. Residues involved in constraining a planar conformation of the ligand are labelled.

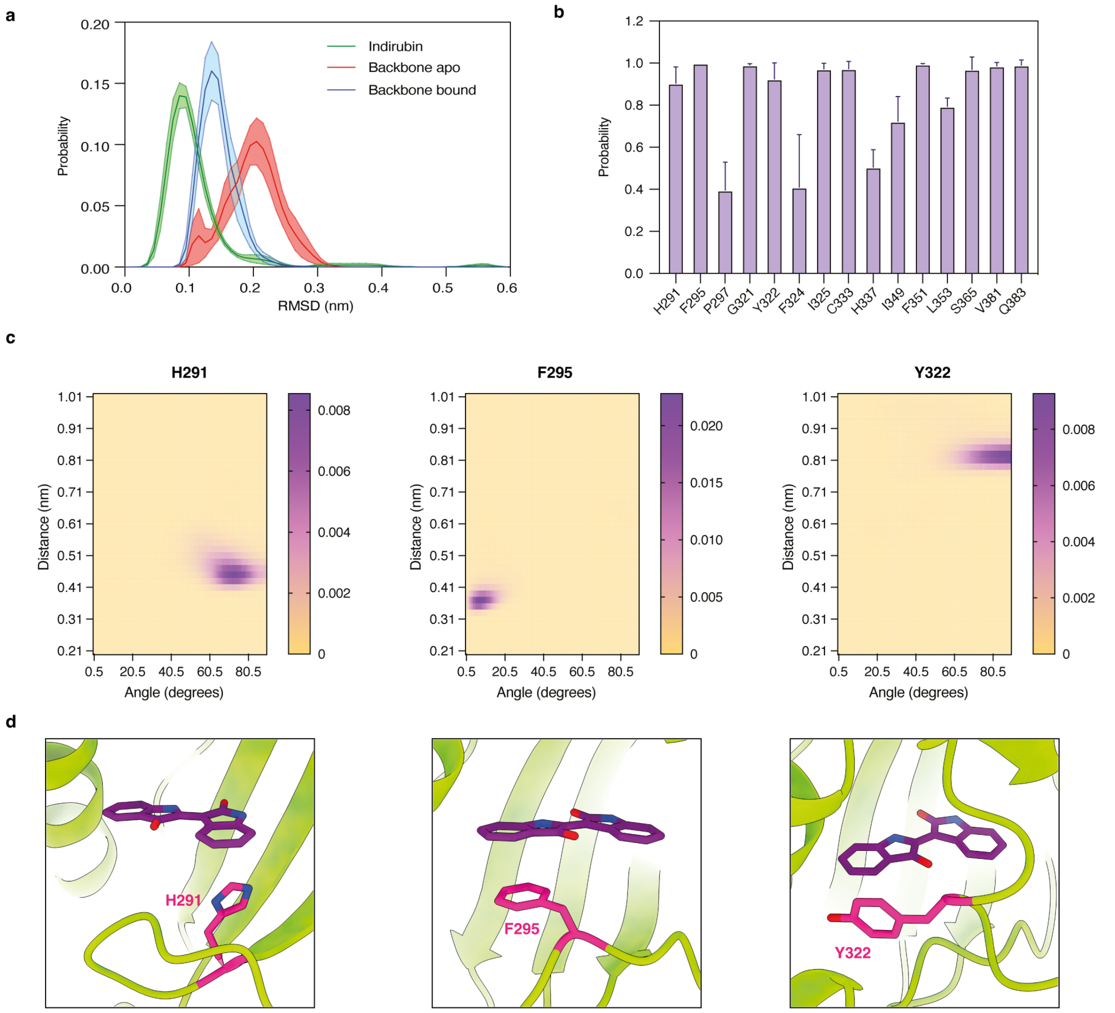


**Extended Data Fig. 9 | Results of the Molecular dynamics simulation for PAS-B domain.** **a**, Probability distribution functions (PDF) of the root mean square deviation (RMSD) of indirubin (green), and of the backbone atoms in the apo (blue) and bound (red) conformations. Values are averages over the MD simulations, while shaded areas are errors of the mean over the replicas. Indirubin remains remarkably stable in the experimental pose in the microsecond timescale, as indicated by an average RMSD with respect to the protein backbone of only 0.11 nm, with limited fluctuations of the position of the ligand. The protein conformation is destabilized by the absence of indirubin, as indicated by larger RMSD of the backbone for the apo system compared to that of the bound system. **b**, Contact probabilities between indirubin and the residues that form contacts with the ligand in the experimental structure. Reported values are averages over the MD simulations, while error bars are standard deviation over the replicas. Most of the contacts observed in the cryo-EM structure are remarkably stable in MD simulations in the microsecond timescale. Other contacts observed in the experimental structure are less stable, but these are formed with residues that are located at distances comparable with the cut-off value used for the definition of contacts. **c**, Two dimensional distributions of the distances and the angles between the aromatic rings of H291, F295, and Y322 with the indole rings of indirubin. (Left) H291 forms a stable T-shape interaction with an average distance between the centers of geometry of the rings of around 0.45 nm and angles between 60 and 80 degrees. (Center) F295, instead, forms a remarkably stable stacked $\pi$-$\pi$ interaction with the indole ring of indirubin, with distances and angles between the two rings that are typical for this type of interaction. (Right) Lastly, Y322 and indirubin assume a T-shape conformation characterized by a rather long distance between the centers of geometry of the two rings, which is nonetheless stable in the microsecond timescale of the MD simulations. **d**, Close-up view of the interactions between residues H291, F295 and Y322 and indirubin.

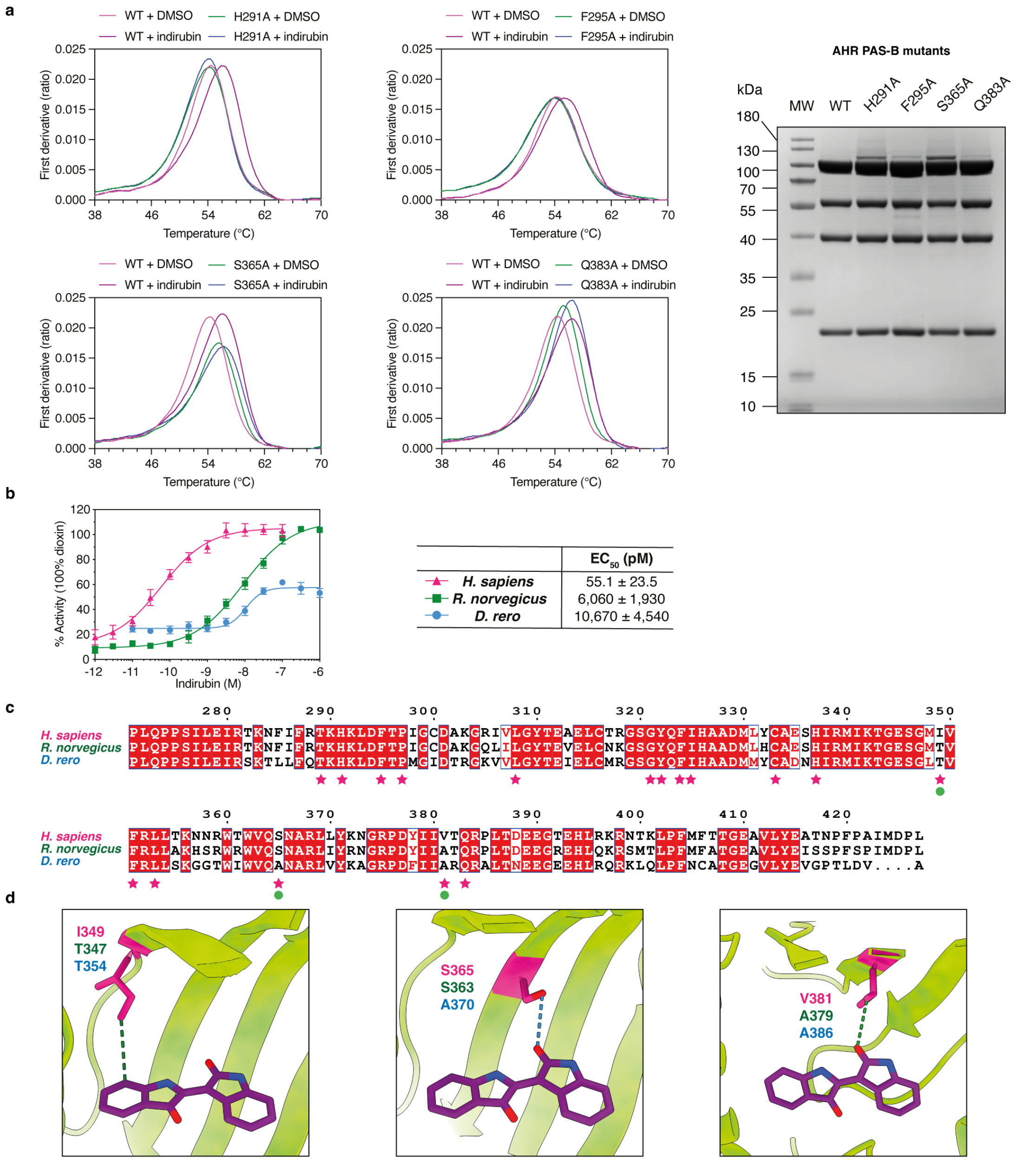


**Extended Data Fig. 10 | Detailed analysis of the indirubin interaction with AHR. a**, Nano-DSF analysis of the indirubin binding to the AHR PAS-B mutants. Mutations H291A, F295A and S365A render protein inactive in the indirubin binding assay. On the other hand, the Q383A mutation does not impair the interaction with indirubin. The first derivative of the ratio between intrinsic fluorescence at 350 nm and 330 nm is plotted against the temperature. The results are representative of independent biological replicates (n=3). The SDS PAGE gel shows the results of the purification for the prepared mutant proteins. The introduced mutations within PAS-B domain did not affect the formation of the complex. 10 μg of protein complex were loaded into each lane. The results are representative of independent biological replicates (n=2). **b**, Indirubin activity in transactivation assays using HAhLH (human), H4AhLH (rat) and ZAhLH (zebra fish) reporter cell lines. Indirubin is a much more potent activator of the human AHR compared with rat or zebra fish orthologs. The values of measured $EC_{50}$ are indicated. For the human AHR, the same plot as in Figure 3B is shown. The results are representative of independent biological replicates (n=4). **c**, Sequence alignment of AHR sequence between three species corresponding to human residues 271-427. Residues that are conserved between all species are highlighted in red. Residues involved in the interactions with indirubin based on the cryo-EM structure are marked with magenta stars. Rat protein differs from human in 2 and zebra fish in 3 of those positions (marked additionally with green circles). **d**, Close-up view of the interactions formed by the residues I349, S365 and V381. Van der Waals interactions are indicated as dashed green lines. Hydrogen bonds are indicated as dashed blue lines.

**Extended Data Table 1 | Cryo-EM data collection, refinement and validation statistics.**

| | **Hsp90-XAP2-AHR with indirubin** (EMDB-14971) (PDB 7ZUB) | **XAP2-AHR** (EMDB-14972) |
|---|---|---|
| **Data collection and Processing** | | |
| Microscope | Titan Krios G3 | |
| Voltage (kV) | 300 | |
| Camera | Gatan K3 | |
| Magnification | 81,000 | |
| Pixel size (Å) | 1.086 | |
| Exposure rate on detector (e–/pix/sec) | 15.0 | |
| Exposure rate on specimen (e–/pix/sec) | 16.5 | |
| Number of frames | 43 | |
| Defocus range (μm) | -2.7 to -1.5 | |
| Automation software | EPU | |
| Energy filter slit width (eV) | 20 | |
| Micrographs collected (no.) | 9,300 | |
| Total extracted particles (no.) | 11,546,649 | |
| **For each reconstruction** | | |
| Final particle images (no.) | 678,724 | 116,399 |
| Point group | C1 | C1 |
| Map resolution (masked) (Å) | 2.76 | 4.00 |
| FSC threshold | 0.143 | 0.143 |
| Map resolution range (Å) | 2.78-5.34 | 3.76-5.26 |
| 3DFSC Sphericity | 0.965 | 0.838 |
| Map sharpening B factor (Å$^2$) | -72.67 | -192.52 |
| Map sharpening methods | RELION3.1.1 | RELION3.1.1 |
| **Model Refinement** | | |
| Initial model used (PDB code) | 5FWP, 4F3L, 4ZPR, 2LKN, 4AIF | |
| Model resolution (Å) | 2.88 | |
| FSC threshold | 0.5 | |
| Model composition | | |
| Non-hydrogen atoms | 28,123 | |
| Protein residues | 1,722 | |
| Ligands | 2 MG, 2 MOO, 1 JY6, 2 ADP | |
| *B* factors (Å$^2$) | | |
| Protein | 64.56 | |
| Ligand | 32.57 | |
| R.m.s. deviations | | |
| Bond lengths (Å) (# > 4σ) | 0.004 (0) | |
| Bond angles (°) (# > 4σ) | 0.946 (1) | |
| Validation | | |
| MolProbity score | 5.44 | |
| Clashscore | 1.62 | |
| Poor rotamers (%) | 1.10 | |
| Ramachandran plot | | |
| Favored (%) | 95.72 | |
| Allowed (%) | 4.28 | |
| Disallowed (%) | 0.00 | |
